# SpineDL: a Deep Learning-based approach for neuron and anatomical structure segmentation in immunofluorescence images of damaged spinal cords

**DOI:** 10.64898/2025.12.17.694828

**Authors:** Pablo Ruiz-Amezcua, Daniel Franco-Barranco, David Reigada, Teresa Muñoz-Galdeano, Rodrigo Martíez Maza, Manuel Nieto-Diaz

## Abstract

In this study, we present SpineDL, an open-source deep learning (DL) approach for neurons and anatomical structure segmentation of the spinal cord in fluorescence images immunostained with NeuN and DAPI, within the context of murine models of spinal cord injury (SCI). SpineDL comprises two main modules: 1) SpineDL-Structure, for semantic segmentation of key spinal cord structures: gray matter, white matter, ependyma, and damaged tissue; and 2) SpineDL-Neuron, for instance-level identification of neuronal somas. To train the models, we developed the SpineDL dataset, a curated collection of 161 confocal images of mouse spinal cord, manually annotated by experts and organized into specific subsets. Both models are based on the HRNetV2-W48 architecture and were trained using state-of-the-art data augmentation and optimization techniques, implemented within the BiaPy framework, following an iterative refinement process driven by quantitative evaluation and expert feedback. Our results show that SpineDL achieves expert-level performance for both cases of structural segmentation and neuron identification. This work provides a robust, reproducible, and extensible platform for the spatial analysis of neurodegeneration following spinal cord injury, representing a step toward the automation of histopathological workflows in neuroscience.

## 1 Introduction

Spinal cord injury (SCI) affects an estimated 15 million people worldwide, leading to permanent motor, sensory, and autonomic deficits with substantial personal, medical, and economic consequences, which is estimated at $1.5–6.2 million per patient in the USA alone [1]. Despite decades of research, no treatment has yet demonstrated sufficient efficacy and security to enter routine clinical use [2].

Neuronal death is a key pathological event driving functional deficits after SCI. In the traumatic SCI, neuronal death occurs in two phases: (1) a primary immediate mechanical focal necrosis and (2) an extended secondary damage of diffuse programmed cell death driven by biochemical and inflammatory events triggered by the trauma [3–5]. This secondary phase encompasses multiple forms of regulated cell death, such as apoptosis, necroptosis, ferroptosis, and pyroptosis [4–6], which might be regulated with appropriate neuroprotective treatments [5]. Understanding this heterogeneity is essential for identifying vulnerable neuronal populations and refining the design of neuroprotective interventions. However, their cellular, spatial and temporal distributions remain largely unknown.

Animal models, particularly rodents, are the primary source of insight into the pathophysiology of SCI, and histology remains the gold standard for analyzing neuronal death and tissue damage. This accumulated body of data constitutes a key resource for understanding the spatial distribution of neuronal death and its relationship to injury characteristics or therapeutic interventions. Accurate identification of neuronal somas is a prerequisite for systematic, spatial analysis of neuronal survival and loss. However, the extraction of such information relies heavily on manual segmentation, which is both time-consuming, error-prone, and difficult to scale for the large datasets required to capture the spatial complexity of injured spinal cord [7].

Over the past decade, deep learning (DL) has driven significant advancements in computer vision applications for the analysis of SCI. DL-based approaches have enabled automated analysis of MRI images in humans [8–11], or SCI classification based on electromyographic data in non-human primates [12]. In murine models, DL techniques have been employed to segment axons and myelin in electronic and light microscopy images [11, 13] or to segment neuronal somas in 3D fluorescence and optic images [14, 15]. However, tools for the automatic segmentation of neurons in the injured spinal cord are lacking and a comprehensive, image-based characterization of neuronal death remains largely unaddressed [7]. Furthermore, the analysis of distinct spinal cord structures is crucial for accurate image registration and 3D reconstruction, as shown in previous studies such as [16], which presents a novel 3D anatomical atlas, and in works focused on the segmentation of gray matter [17] and white matter [11, 13].

In this study, we present SpineDL, an open-source deep learning-based tool designed for the segmentation of spinal cord neurons using fluorescence images immunostained against Rbfox3 (usually known as NeuN, the most commonly used marker of postmitotic CNS neurons), and DAPI (4’,6-diamidino-2-phenylindole, a common fluorescent stain for nuclei) in the context of SCI. The tool has been developed using BiaPy [18], an open-source library tailored for constructing deep learning workflows in bioimage analysis. SpineDL offers two main modules: 1) SpineDL-Structure, for automatic segmentation of key anatomical structures of the spinal cord, including the gray matter, white matter, ependyma, and damaged regions; and 2) SpineDL-Neuron, for automatic identification of neuron somas.

In parallel with the development of the tool, we introduce the SpineDL dataset, a curated collection of confocal fluorescence images of mouse spinal cord sections labeled with NeuN and DAPI. The dataset compiles images from two previously published studies [19, 20] employing murine models of traumatic SCI and includes comprehensive manual annotations for both spinal cord structures and neuronal somas. These annotations were generated through iterative refinement involving deep neural network feedback and expert review, resulting in two high-quality subsets: one for structure segmentation and another for neuron identification. The final dataset, comprising 161 annotated images across training, validation, and test sets, is publicly available through Zenodo, providing a valuable resource for benchmarking and advancing deep learning approaches in spinal cord histopathology.

The aim of SpineDL is to provide an accessible, flexible, and reproducible platform for sptial analysis of neuronal death in SCI models. The tool enables automated segmentation of spinal cord structures and neurons before and after injury. This workflow facilitates the study of injury-associated anatomical changes and offers a robust framework for investigating cellular mechanisms underlying SCI pathophysiology.

We summarize our main contributions as follows:

**–** Introduces SpineDL, a novel open-source deep learning-based tool designed specifically for automated segmentation and analysis of spinal cord fluorescence images in the context of spinal cord injury.
**–** Release of the SpineDL dataset, a publicly available, expertly annotated collection of spinal cord fluorescence images from murine SCI models, comprising separate subsets for structure and neuron identification. The dataset provides a standardized benchmark for developing and validating deep learning methods in spinal cord histology.

## 2 Methods

### 2.1 SpineDL dataset Dataset adquisition

The images analyzed in this study were sourced from two independent, previously published investigations employing mouse models of traumatic spinal cord injury. The first, [20], examined cell-specific alterations in autophagy following contusive SCI. The second, [19], evaluated the neuro-protective effects of the anti-apoptotic drug UCF-101. In both studies, confocal microscopy images were acquired from 20*µm* transverse sections of the thoracic spinal cord of adult mice (Mus musculus, strain C57BL/6). Sections were fluorescently stained with two routine markers: an antibody against NeuN to label neuronal somas and DAPI as a general nuclear counterstain.

Full spinal cord sections were imaged as mosaics of tiled images using either a High-Speed Resonant Confocal Scanner Microscope or a TCS SP5 fast-scan confocal microscope (Leica Microsystems CMS GmbH, Wetzlar, Germany), both equipped with PL APO 20×/0.70 CS dry UV objectives. The original raw images were acquired as Z-stacks at varying resolutions from 0.445 to 0.891*µm*/pixel with 8 to 32-bit depth and stored in .lif files. Preprocessing included maximum intensity projection of the stacks, the arrangement of the images into two grayscale channels (Channel 1: DAPI; channel 2: NeuN) and their conversion to TIFF format. Additionally, some images were cropped for training on specific features. All preprocessing was carried out using Fiji. Images were split into training, validation and testing subsets, care was taken to balance the number of images from each source in the subsets. Images and metadata are accessible in our Zenodo repository.

### Dataset annotation

#### Structure identification dataset

The dataset was developed through an iterative process, as illustrated in Figure 1. To reach its final form, a total of eight iterations were conducted. The first iteration served as a preliminary step and a proof of concept to evaluate whether a deep neural network (DNN) could effectively address the problem. In this initial phase, ten images were manually labeled with only two classes: gray matter and background.

**Fig. 1:**
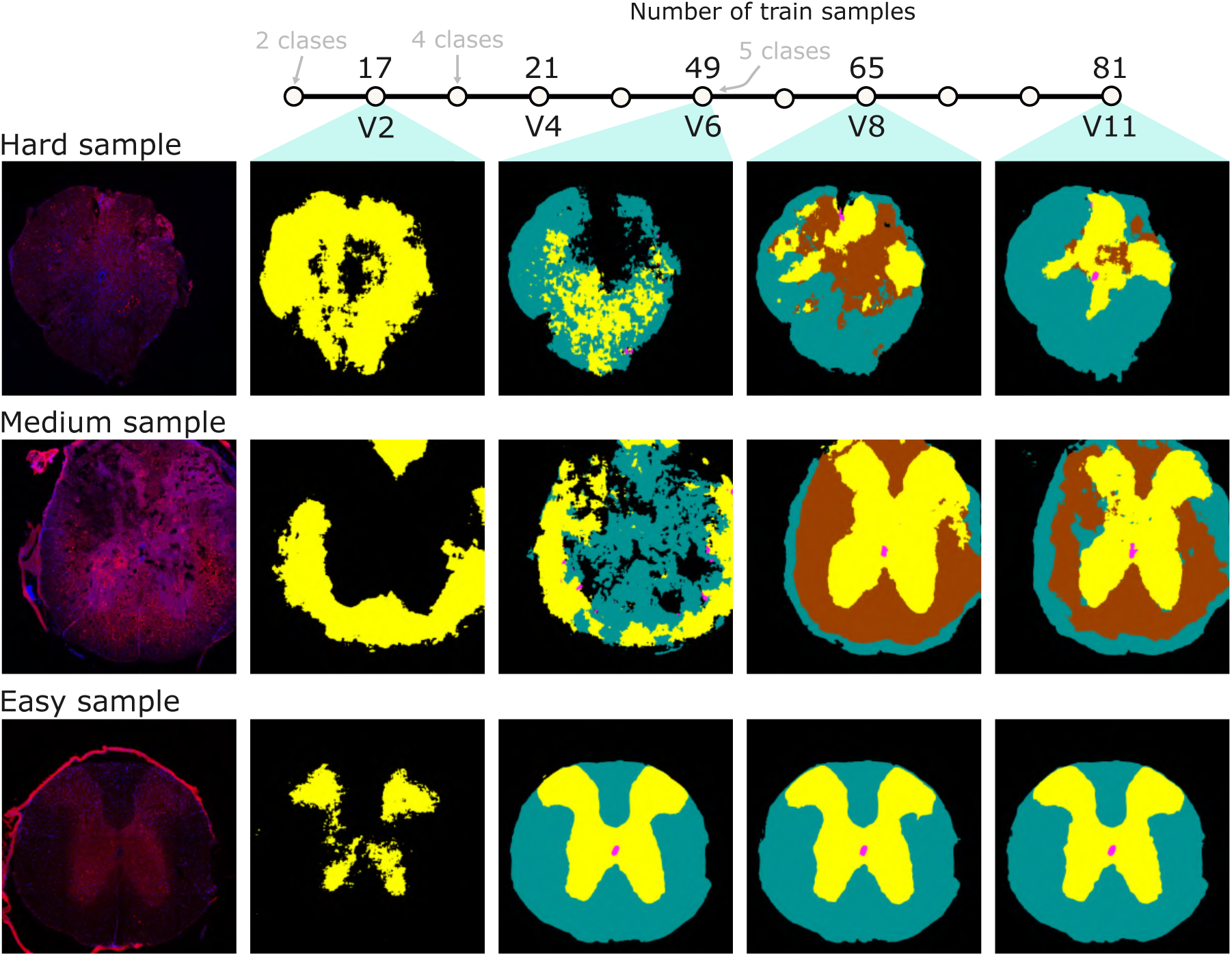
Overview of the process used to construct the structure identification dataset. This process underwent eleven iterations. The first column displays selected raw images from each dataset, and the remaining columns show the semantic segmentations produced by the DNN across the different iterations. The process involved refining the labels generated by the DNN in previous iterations and gradually increasing the number of training samples. Notice that for the creation of this data new classes were also added during the iterative process. Each dataset includes three representative examples categorized as easy, medium, and hard, based on the level of difficulty the DNN encountered in generating accurate segmentations. As illustrated, the DNN’s performance improves steadily across iterations, ultimately achieving what we consider a low error rate.

Following promising results, additional training images were incorporated. Some were manually labeled from scratch, while others were labeled using predictions generated by the DNN itself and curated afterwards, thereby accelerating the annotation process. In the third iteration, once the DNN demonstrated acceptable performance in predicting gray matter, two new classes were introduced, resulting in a total of four: white matter, gray matter, ependyma, and background. While the DNN performed well on some images, others remained challenging.

To improve performance, the DNN was used to predict labels on new images, and those with poor prediction quality were selectively added to the training set. This strategy of targeted data addition was repeated in subsequent iterations, progressively expanding the training set and enhancing DNN accuracy.

By the sixth iteration, four tissue classes were being predicted with satisfactory performance. At this point, the most complex class was added: the injured tissue. By then, the training set included 49 images, and the DNN performed reliably on images from non-injured sections. The final two iterations focused on refining labels and adding more images with injured tissue to the training set. Identifying injured regions proved particularly difficult, not only for the DNN but also for human experts, as the staining was not aimed at identifying damaged tissue but neurons and cell nuclei. Accurate annotation of these regions posed a significant challenge, but it was essential for ensuring sufficient data quality and quantity to train the neural network effectively. As additional samples were incorporated and existing annotations refined through successive iterations, the final version of the dataset, corresponding to the eleventh iteration, comprises 81 carefully annotated images (74 for training and 7 for validation), encompassing the five target classes: white matter, gray matter, ependyma, injured tissue, and background.

For the test set, a group of 3 independent expert annotators analyzed a set of 10 images, manually delineating the boundaries of the spinal cord section, excluding the dorsal root ganglia, the meninges and any obvious artifacts, as well as the gray matter, the ependyma, and the damaged tissue.

The evolution of the dataset necessitated a corresponding adjustment and refinement of the underlying DNN architectures and training configurations. Across each iteration, multiple DNN configurations were rigorously tested to maximize performance metrics and, critically, to minimize the time required for expert curation and proofreading of the machine-generated labels on new samples.

The specific hyperparameters and architectural choices explored throughout the iterative development process are thoroughly documented in the supplementary material (Table S1). The model configuration that ultimately demonstrated performance metrics comparable to, or exceeding, inter-expert agreement, specifically, the final configuration trained on the 81 images, eleventh iteration dataset, is formally designated as SpineDL-Structure. This final model represents the culmination of the data curation effort, delivering a robust solution capable of accurate, five-class structural segmentation across the delineated spinal cord anatomy.

#### Neuron identification dataset

Similar to the structure identification dataset, this dataset was constructed through an iterative process, gradually incorporating more training images and refining the segmentation masks generated in previous iterations. The new training images in each iteration were selected by identifying those images and regions with low segmentation accuracy, i.e. where major artifacts or erroneous segmentations were present. In this case, seven iterations were performed, resulting in a final set of 82 (75 for training and 7 for validation). The complete process is illustrated in Figure 2.

**Fig. 2:**
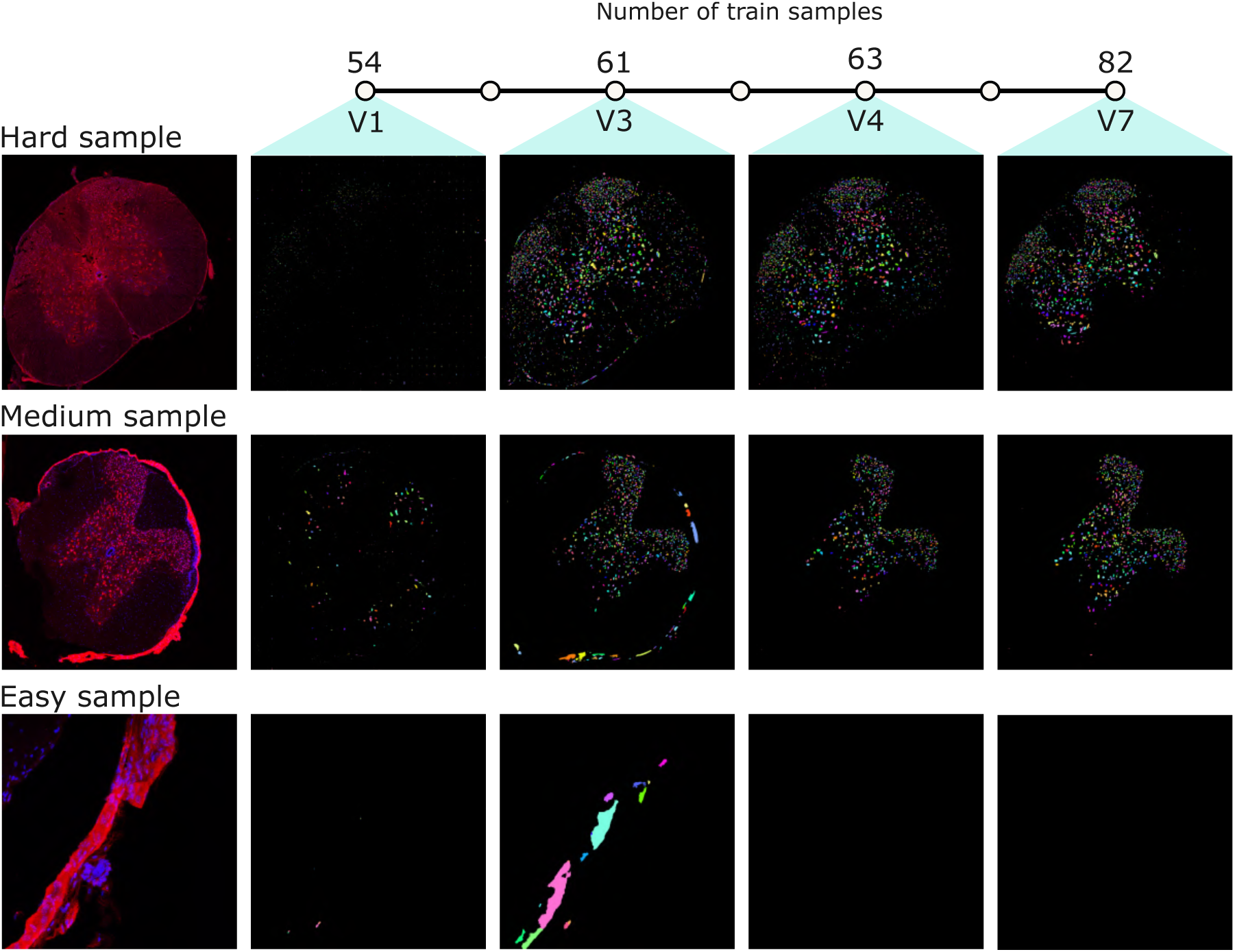
Overview of the process used to construct the neuron identification dataset. This process underwent seven iterations. The first column displays selected raw images from each dataset, and the remaining columns show the segmentations produced by the DNN across the different iterations. The process involved refining the labels generated by the DNN in previous iterations and gradually increasing the number of training samples. Each dataset includes three representative examples categorized as easy, medium, and hard, based on the level of difficulty the DNN encountered in generating accurate segmentations. As illustrated, the DNN’s performance improves steadily across iterations, ultimately achieving what we consider a low error rate.

A particular challenge encountered during the development of this dataset was the DNN’s tendency to incorrectly identify neurons in regions affected by staining artifacts, such as the meninges depicted in the “medium sample” of Figure 2, which does not contain neuronal nuclei. To address this issue, image patches containing such regions were added to the training set, enabling the DNN to learn to distinguish these areas as non-neuronal.

For the test set we employed two sets of images. The first one, composed of 5 images from [19], was the test set of a benchmark designed to compare the precision, repeatability, and reproducibility of manual, threshold-based, and DL-based neuron identifications [7]. In this study, we added a second set of 5 images from [20] to complete the test set including different tissue processing, staining and imaging conditions. Images were selected to cover the different conditions of the sections, from uninjured spinal cords with clear neuron stainings to injured cords with strong background. Each image of these test sets was independently annotated five times by three expert annotators (two of whom performed replicate analyses, and one conducted an additional analysis). During each session, the expert annotators marked all objects they identified as neurons. The annotations were merged, and each candidate object was assigned a likelihood of being a neuron, defined as the number of times it was identified divided by the total number of annotations for that image (i.e., five).

As with the structural analysis model, the development of the neuron segmentation approach involved continuous adaptation of the underlying DNN architecture across the seven iterations. The goal remained to achieve performance parity with, and ultimately surpass, expert-level agreement in identifying individual neuronal nuclei. The configurations and hyperparameters explored throughout this parallel development effort are cataloged in the supplementary material (Table S2). The final model configuration, trained on the comprehensively refined 82-image dataset (the seventh iteration), and optimized to accurately manage the complexities introduced by artifacts and diverse staining conditions, is designated as SpineDL-Neuron.

### 2.2 Custom DNN architectures: 2D HRNetV2

Both DNNs, the one for segmenting anatomical structures (SpineDL-Structure) and the other for neuron identification (SpineDL-Neuron) follow the High-Resolution Network: HRNetV2-W48 [21]. The DNN is a 2D convolutional architecture designed to preserve high-resolution representations throughout the entire feature extraction process. Unlike traditional encoder-decoder architectures that recover spatial details after successive downsampling, HRNet maintains multiple parallel branches operating at different resolutions and continuously exchanges information across them. This design enables the network to capture both fine-grained spatial information and robust contextual features. The W48 variant uses 48 channels in the high-resolution branch and progressively adds lower-resolution branches with increased channel capacities. Batch normalization and ReLU activations are applied after each convolutional layer to improve training stability and regularization.

Both models are fully integrated within the BiaPy library [18], using Pytorch version 2.9.1. We used a patch size of 512 *×* 512 in both models. We employed data augmentation techniques such as, horizontal and vertical flips, brightness and contrast changes, elastic transformations and zoom variations. Cross entropy was used as a loss function, with a learning rate of 1*e −* 4, employing a cosine-decay scheduler with warm-up [22], and ADAMW optimizer [23]. Both DNNs were trained until convergence using four NVIDIA GeForce RTX 3090.

Following the code of good practices to show deep learning-based results proposed by [24], a thorough hyperparameter exploration throughout the SpineDL dataset creation of both models presented here, i.e. model SpineDL-Structure and SpineDL-Neuron, can be found in Table S1 and Table S2 respectively. Each DNN follows a specific configuration tailored for each of the tasks as is described as follows:

#### SpineDL-Structure for anatomical structure segmentation

Specifically designed for a semantic segmentation task, this DNN accurately segments five classes corresponding to key anatomical structures of the spinal cord, including the background, gray matter, white matter, ependyma, and damaged regions such as lesions or imaging artifacts.

#### SpineDL-Neuron for neuron identification

To specifically address the task of neuron identification, i.e. an instance segmentation task, we employed a bottom-up approach inspired by the method described in [25]. The DNN architecture is trained to simultaneously predict two outputs: a binary foreground segmentation mask (F) and an instance center of mass (P), which can be referred to as ‘FP’.

After prediction, both outputs are thresholded and combined to enhance object delineation. A connected components algorithm is then applied to the combined mask to identify discrete, non-overlapping neuron instance seeds. These seeds are subsequently used as markers in a marker-controlled watershed algorithm [26], which incorporates three key components: (1) the inverted fore-ground probability map as the input image (defining the topographic surface), (2) the instance seeds as the marker image (indicating the initiation points of the flooding process), and (3) a binarized version of the foreground mask (B) as the constraint mask (limiting the extent of region expansion). This integrated pipeline enables the effective segmentation of individual neuron instances. A visual representation of the process is provided in Figure 3.

**Fig. 3:**
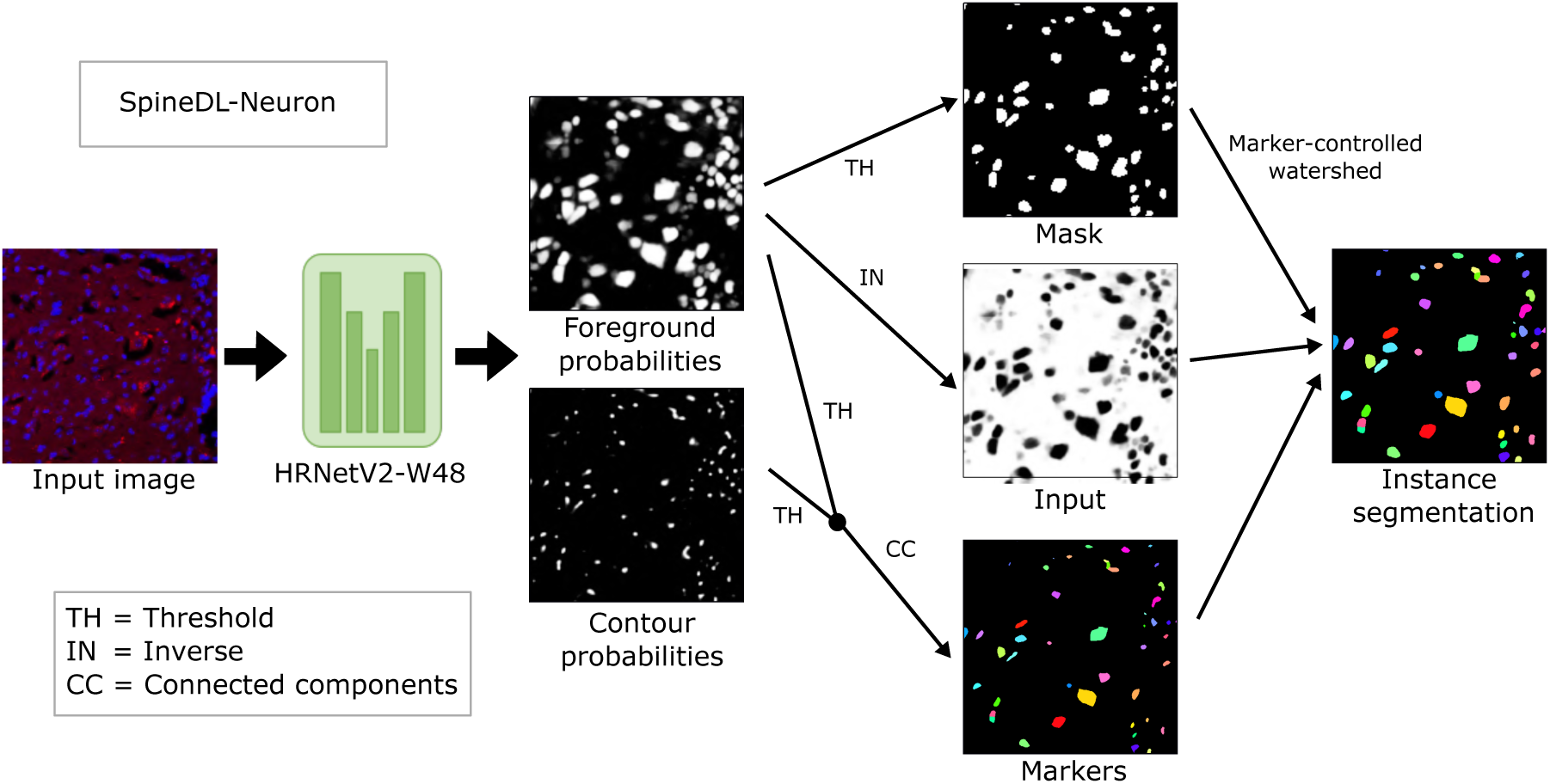
Processing pipeline of our proposed DNN for neuron identification. The DNN predicts foreground and center of mass probabilities that are fused to create three inputs for a marker-controlled watershed [26] to produce individual instances.

### 2.3 Evaluation metrics

Given that the two DNNs address distinct tasks, semantic segmentation in the case of SpineDL-Structure, and instance segmentation for SpineDL-Neuron, the evaluation protocols also differ accordingly.

#### SpineDL-Structure for anatomical structure segmentation

We used the widely adopted Intersection over Union (IoU), which measures the overlap between two instances (A, B) and can be calculated as:

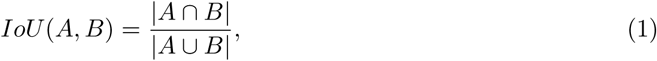

being *∩* the intersection and the *∪* union.

#### SpineDL-Neuron for neuron identification

For each image, the five manual annotation sessions were aggregated and expert clicks were grouped into putative neurons using DBSCAN (*ɛ* = 8*px*) [27]. This procedure does not alter any ground truth (GT) coordinates; it serves only to merge clicks from different experts that refer to the same anatomical object within a small spatial tolerance. Each resulting “manual object” was assigned a class equal to the number of distinct experts who clicked it (1–5).

To introduce a tolerance in matching between predicted instances and GT clicks, every predicted instance was dilated with a disk structuring element of radius 8 px, after which point-in-mask membership was tested against the dilated mask. A manual object was deemed detected by the DNN if any of its constituent clicks fell inside any dilated predicted instance for that image; multiple overlaps did not increase the count for that object. DNN-only detections (class 0) were defined as predicted instances whose dilated mask contained no manual clicks from any session.

For each test image, two distributions over classes 0–5 were computed: (i) Manual, the counts of manual objects in classes 1–5 (class 0 = 0 by definition); and (ii) DNN, the counts of DNN detections for manual objects in classes 1–5, with class 0 capturing DNN-only instances.

### 2.4 Community-Oriented Tools and Resources

To ensure that SpineDL DNNs are accessible and reusable by the scientific community, and in line with the best practices advocated by BiaPy [18], two comprehensive tutorials have been developed for each workflow, i.e. SpineDL-Structure and SpineDL-Neuron detailing how the DNNs can be reused in other works.

In addition, as BiaPy is a community partner of the BioImage Model Zoo [28], the DNNs have also been published on that platform, enabling seamless integration with other tools and facilitating their rapid adoption in diverse bioimage analysis applications. Specifically, SpineDL-Structure model can be found as ‘greedy-deer’ and SpineDL-Neuron as ‘proactive-snail’.

## 3 Results

### 3.1 Structure segmentation results

The agreement matrices in Figure 4 provide a detailed overall and per-class IoU scores between the DNN predictions and each expert annotation, averaged over all test images, a metric referred to as the mean IoU (mIoU). Overall, the model achieves an average IoU of around 0.7, comparable to the consistency observed among the human experts themselves. The lowest expert–expert agreement is 0.69 (Expert 3 vs Expert 2) and the highest is 0.86 (Expert 1 vs Expert 3), defining the realistic lower and upper bounds of human consistency for this task. Since the model’s performance falls squarely within this human range, it can be interpreted as behaving like an additional expert annotator. This range also underlines the inherent difficulty of the segmentation problem: even among specialists, perfect consensus is rare.

**Fig. 4:**
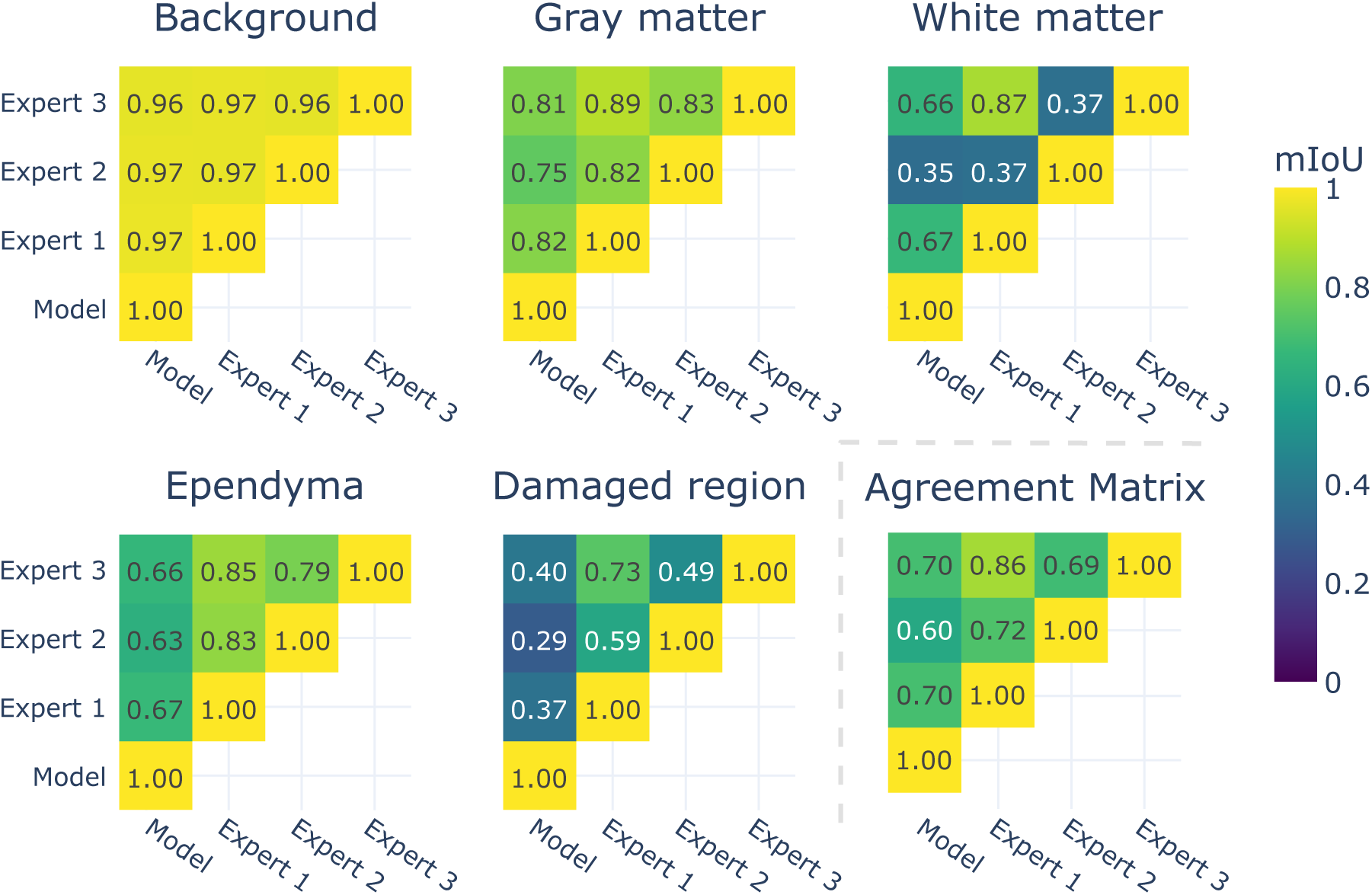
Agreement Matrices (Dataset-Level IoU). **A-E.** Per-class IoU agreement matrices between the DNN and all expert annotations. High agreement is observed for structured regions (gray and white matter), while lower scores in the damaged region reflect higher annotation uncertainty. **F.** Overall pairwise IoU among the DNN and experts. The model-expert agreement (*≈* 0.70) matches inter-expert consistency, ranging from 0.69 to 0.86, indicating the DNN performs comparably to a human annotator.

At the class level, the results confirm that segmentation difficulty varies strongly across structures. High agreement is observed for more structurally defined regions such as the spinal cord section (as opposed to background), the gray matter, and the ependyma, all with consistent boundaries and clear visual definition across annotators and the model. In contrast, the damaged region class shows markedly lower IoU values, even among human experts, highlighting its subjective and irregular nature. Intermediate values are observed in the white matter, whose distinction from damaged tissue is ambiguous in injured sections. This pattern highlights how both the model and experts find well-defined anatomical structures easier to segment reliably, whereas irregular or ill-defined boundaries remain challenging even for trained annotators. This reinforces that discrepancies in this class primarily stem from annotation uncertainty rather than model limitations, illustrating the intrinsic challenge of delineating pathologic or irregular areas.

Additional analyses providing class-level agreement details and model performance against the consensus ground truth further illustrate the variability of agreement across individual tissue classes and the model’s behavior relative to an expert consensus reference. As shown in Figure 5, the IoU obtained when comparing the model’s predictions with the unanimous-consensus ground truth, a reference mask generated by merging all expert annotations and retaining only the pixels where full agreement was reached, the model achieves an mIoU of approximately 0.69, with strong performance in gray matter, white matter, and ependyma (0.89, 0.78, and 0.74, respectively) and lower values in the damaged region (0.34), consistent with its greater annotation variability. These results confirm that, when evaluated against the most reliable subset of human agreement, the DNN is highly consistent with expert consensus, approaching the upper range of human-level performance. A qualitative comparison is depicted in Figure 5B.

**Fig. 5:**
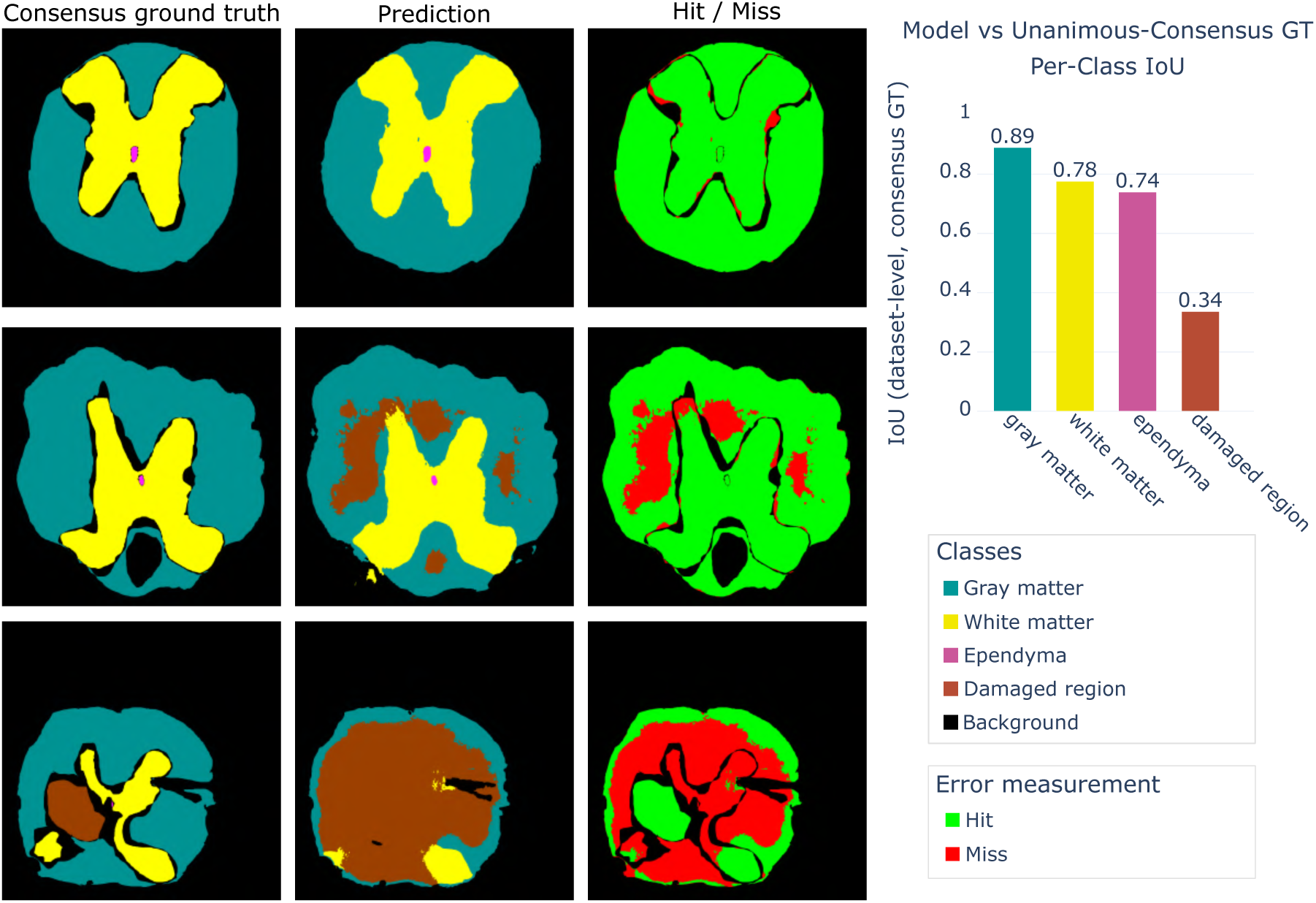
Model vs Unanimous-Consensus GT. **A:** Qualitative Comparison. Examples showing unanimous-consensus ground truth (left), the DNN predictions (middle), and pixelwise agreement maps (right) in three images of differing difficulty. The visualization illustrates that most discrepancies occur along structure boundaries and within damaged regions, consistent with their higher annotation uncertainty. **B:** Per-class IoU between the DNN predictions and the unanimous-consensus ground truth, built from pixels where all experts agreed. The model shows expert-level performance on well-defined classes and lower agreement on ambiguous damaged areas.

### 3.2 Neuron segmentation results

The confidence threshold used to define neuron seeds in the center-of-mass (P) channel, i.e., to determine where candidate instance centers are selected in the bottom-up segmentation pipeline (as shown in Figure 3), was empirically optimized using the validation dataset through a precision-recall (PR) analysis (see Supplementary Figure S1). This analysis, based on an instance-level evaluation (described in Supplementary), revealed that thresholds in the range of 0.3 *−* 0.4 provided the most balanced trade-off between precision and recall. Accordingly, we fixed the seed-selection threshold at 0.3 and set the foreground constraint threshold that limits region growth during the watershed step to 0.9 after qualitative inspection of the predictions.

Figure 6 summarizes the agreement between manual and DNN identifications across consensus classes. The ‘Manual’ series reflects the distribution of manual objects by how many of the five sessions identified them (classes 1–5; class 0 = 0 by definition). The DNN series reports, for each class, the frequency with which the DNN detected those manual objects, together with class 0 capturing DNN-only instances (no manual clicks within the 8-px tolerance introduced via dilation). Because values are normalized per image and averaged across images shared by the GT and prediction sets, the bars provide a scale-free view of (i) how detections vary with manual consensus and (ii) the prevalence of algorithm-only detections. This visualization complements thresholded, one-number metrics by revealing class-specific behavior that is otherwise obscured.

**Fig. 6:**
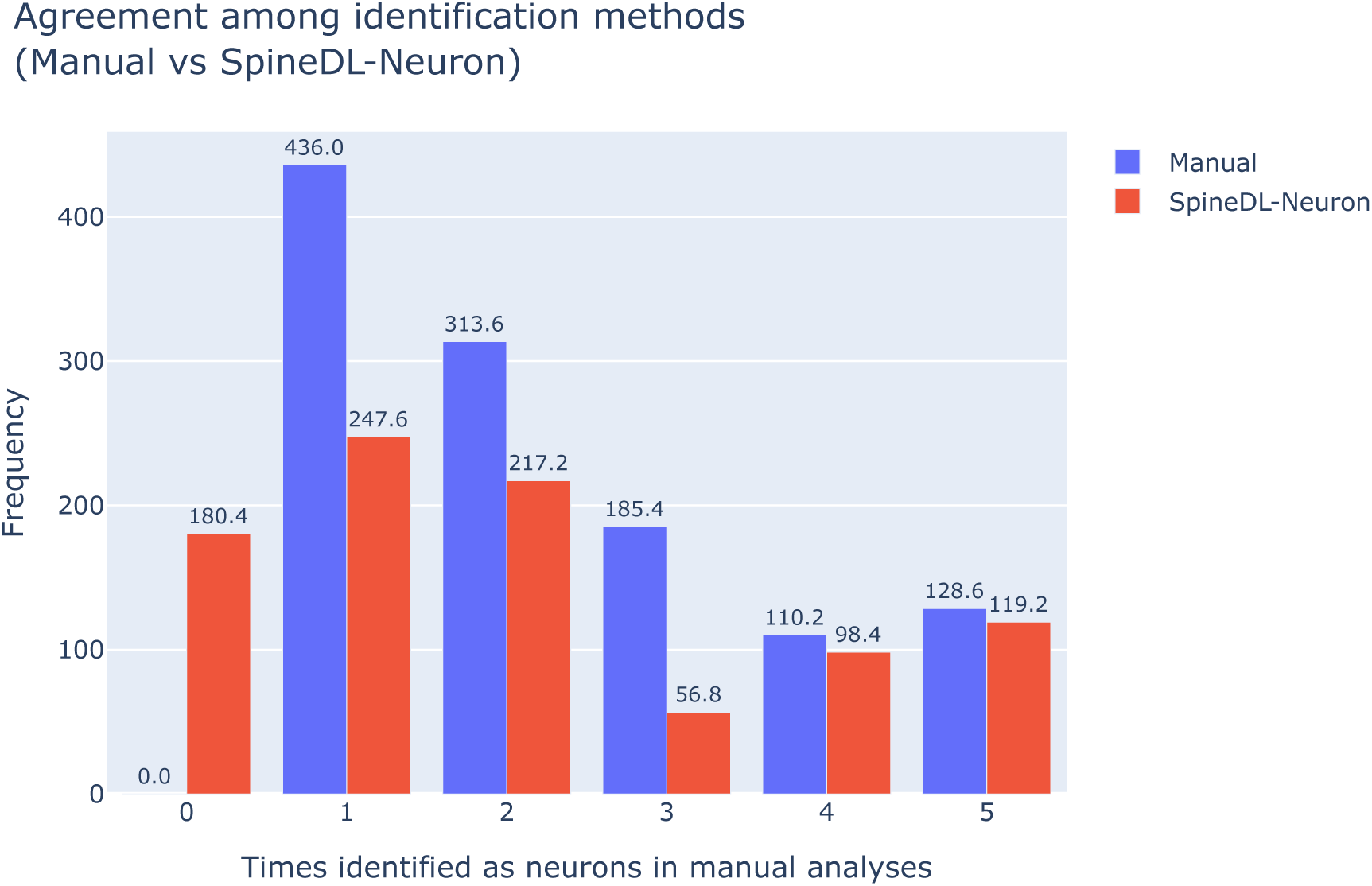
Agreement between manual and DNN identifications with tolerance-aware matching. Grouped bars show Manual and DNN series for classes 0–5, averaged across images present in both the ground-truth and prediction sets. Class numbers (1–5) indicate how many of the five manual sessions identified an object as a neuron; class 0 denotes DNN-only detections (no manual clicks within an 8 px tolerance). The Manual series reflects the distribution of manual objects across classes 1–5 (class 0 = 0), whereas the DNN series reports, for each class, the proportion of manual objects detected by the DNN plus the number of DNN-only instances at class 0.

## 4 Discussion

In a previous study [7], we demonstrated that a trained neural network can outperform manual and threshold-based identification methods in segmenting neuron somas from undamaged and damaged spinal cord sections stained with the neuronal marker NeuN and the nuclear marker DAPI. The network, developed using the Olympus software CellSense Dimensions, produced predictions highly consistent with manual annotations, overcoming limitations of reproducibility, repeatability, and time: one of the main constraints in large-scale image analysis. This tool enabled the rapid and efficient assessment of neuronal distribution across dozens of sections from multiple control and SCI individuals with or without antiapoptotic drug treatment.

From these results, two main directions for further development were identified: (1) the need for an open, non-proprietary neural network for neuronal segmentation; and (2) the improvement of image registration systems toward fully automated workflows. The present study addresses both objectives. First, we developed SpineDL-Neuron, an open-source, scalable, and fully documented deep learning model for instance segmentation of neurons, designed to comply with FAIR principles and open science standards. Second, we generated SpineDL-Structure, a second DL model for semantic segmentation of the main anatomical structures of the spinal cord, aimed at facilitating image registration.

To the best of our knowledge, the SpineDL-Structure model represents the first of its kind trained on fluorescence images of injured spinal cords. Its development posed particular challenges, as the staining protocol used (immunofluorescence against NeuN plus DAPI nuclear staining) was designed for neuron nuclear labeling rather than for distinguishing gray and white matter or identifying damaged areas. Despite this, the model proved highly effective, achieving performance comparable to that of histologists specialized in spinal cord injury. Although the overall agreement with the ground truth was moderate (IoU *≈* 0.7), this value was largely influenced by the lower accuracy in detecting damaged tissue areas (IoU *≈* 0.34) as well as by minor discrepancies in boundaries, suggesting that disagreements were largely due to annotation uncertainty rather than limitations of the model. In contrast, predictions for background (and therefore of the section perimeter), residual gray matter, and ependymal regions showed high consistency with experts when considering the areas of consensus among manual annotators. Accurate identification of these regions is especially relevant for image registration, as they provide homologous reference points between target and reference sections. Nevertheless, the delineation of lesion areas remains an important aspect for future model improvements.

The SpineDL-Neuron module employs a bottom-up instance segmentation approach based on center-of-mass detection and marker-controlled watershed, allowing reliable identification of individual neuronal somas similar to those obtained in the previous model developed with Olympus CellSense Dimensions [7]. Direct comparison between the two models, using the same image set defined in [7], allowed for an objective assessment of their relative performance. Both models exhibited a low number of artifacts (defined as predictions not corresponding to neurons identified by at least one expert; see Figure 7), although the present model still showed some susceptibility to artifacts in the lesion zones. These can, however, be efficiently removed when combined with the anatomical segmentation model, as they do not correspond to gray matter regions. The current model also showed some difficulties in identifying large motoneurons, which were correctly detected by the previous model in nearly all cases. Analysis of the segmentation outputs suggests that these discrepancies are more related to the center-of-mass threshold values used to generate the segmentations than to the foreground predictions of the neural network. Adjusting these parameters can correct most of these issues, albeit at the cost of a slight increase in artifact frequency.

**Fig. 7:**
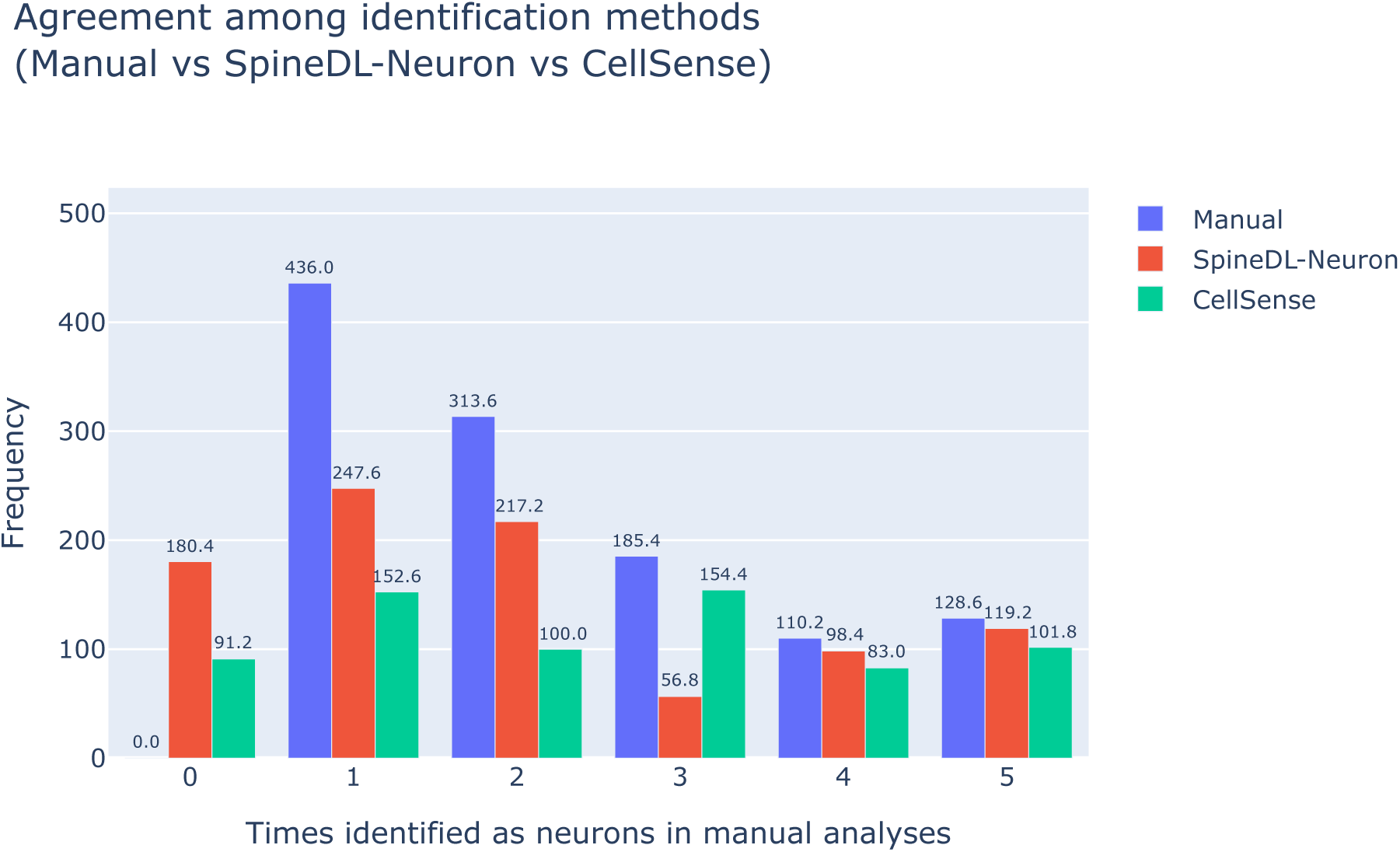
Agreement between manual and DNN identifications with tolerance-aware matching. Grouped bars show Manual, SpineDL-Neuron, and CellSense Dimensions series for classes 0–5, averaged across images present in all three sets. Class numbers (1–5) indicate how many of the five manual sessions identified an object as a neuron; class 0 denotes DNN-only detections (no manual clicks within an 8 px tolerance). The Manual series reflects the distribution of manual objects across classes 1–5 (class 0 = 0), whereas the SpineDL-Neuron and CellSense series report, for each class (1–5), the proportion of manual objects detected by that DNN plus the number of DNN-only in-stances at class 0.

Despite these limitations, the proposed model offers significant advantages over the previous one: 1) It is open source, not a commercial closed software as Olympus CellSense Dimensions, and follows FAIR (Findable, Accessible, Interoperable, and Reusable) principles [29]; 2), both the anatomical and neuronal segmentation models can be retrained by researchers to adapt them to their specific experimental conditions; and 3) SpineDL-Neuron employs an instance segmentation approach to capture individual neuronal nuclei, while CellSense Dimensions generates binary semantic segmentation maps that necessitate subsequent post-processing steps utilizing prior knowledge of the target structure such as object size.

The development of SpineDL is aligned with open science and FAIR principles [29]. The public release of code, trained models, and expert-validated annotations via open open platforms enhances transparency and promotes widespread adoption, validation, and extension of the framework by the scientific community. Moreover, the pipeline is implemented within the BiaPy framework and is fully reproducible, with models and tutorials made available to promote transparency, adaptability, and scalability for the broader scientific community. In addition, the public release of the SpineDL dataset, a curated and expertly annotated collection of confocal spinal cord images, provides a solid foundation for further developments in computational histopathology applied to SCI.

Overall, this work represents a significant advance toward fully automated workflows for histological image analysis, enabling large-scale, systematic studies of neuronal degeneration after SCI.

Several lines of future development can further enhance the applicability and impact of SpineDL. Although the present work focuses on mouse spinal cord, testing the models in rat sections, a widely used and translationally relevant SCI model, will allow us to evaluate the robustness of the segmentation pipeline under differences in tissue size, cytoarchitecture, staining patterns, and injury morphologies. Such validation will help determine the generalizability of SpineDL across species and experimental conditions, and may ultimately support its extension to other animal models or human post-mortem samples.

A second major direction involves the development of a 3D reconstruction pipeline for the spinal cord. The anatomical segmentations produced by SpineDL-Structure will serve as anchors for accurate image registration and integration into 3D anatomical atlases, such as the one developed by [16] for volumetric mapping of neuronal survival. Coupling SpineDL-derived segmentations with these reconstruction methods would enable the generation of spatially normalized representations of neurodegeneration following SCI, thereby facilitating analyses of its relationship with gray and white matter alterations and injury progression. Such 3D reconstructions may offer unprecedented opportunities to study injury propagation, compare injury severity across individuals, and correlate histopathological features with functional outcomes.

## Acknowledgements

We are very grateful to Drs. David Reigada, Teresa Muñoz Galdeano, Hugo Vara, Javier Mazaŕıo, Virginia Vila, Maŕıa Asuncíon de la Barreda Manso, Veŕonica Barranco and Irene Novillo for manual annotation of the test images employed in this study. We are also thankful to the Fundacíon del Hospital Nacional de Parapĺejicos para la Investigacíon y la Integracíon (FUHNPAIIN) for their technical and logistic support. Special thanks to Jose Ángel Rodŕıguez-Alfaro and Javier Mazaŕıo from the Microscopy Facility of the Experimental Neurology Unit (Hospital Nacional de Parapĺejicos, Toledo) for their help in image adquisition and procesing.

## 6 Data availability

SpineDL dataset is available in our Zenodo repository under the CC BY 4.0 license. Additional metadata, raw images, and information on the subjects included in this study are available in the NeuroCluedo OSF repository, also under the CC BY 4.0 license.

## 7 Code availability

The experiments were performed using the BiaPy library [18], an open-source tool whose source code is publicly available on BiaPy’s GitHub. Furthermore, to facilitate the utilization and reproduction of our work, we have developed two comprehensive tutorials: one for the SpineDL-Structure workflow and another for the SpineDL-Neuron workflow. On top of that, both models have been uploaded to the BioImage Model Zoo so users can use them for their work in a more accessible way. SpineDL-Structure model can be found as ‘greedy-deer’ and SpineDL-Neuron as ‘proactive-snail’.

## 8 Author contributions

P.R.A, D.F.B and M.N.D. conceived and designed the work. D.F.B designed the deep learning experiments, while P.R.A, D.R. and M.N.D curated SpineDL dataset. P.R.A, D.F.B and M.N.D. wrote the paper with input from T.M.G and R.M.M.

