## Supplementary Materials for "SpineDL: a Deep Learning-based approach for neuron and anatomical structure segmentation in immunofluorescence images of damaged spinal cords"

### 1 Hyperparameter search space for each configuration

**Notations:**

- $[a, b]$ : Range between two possible values. E.g. $zoom([0.75, 1.25])$ corresponds to a random zoom value between 0.75 and 1.25.
- $[a, b, c]$ : All values from a to b with c step. E.g. $[10, 300, 10]$ corresponds to $10, 20, 30, 40, ..., 300$
- $choice[a, b, ...]$ : one value between a, b and so on. E.g. $[10, 15, 20, 30, 60]$ possible values are: 10 or 15 or 20 or 30 or 60 (but only one).
- $a, b, c, ...$ : all tested values. E.g. flips, rotations, etc.

The hyperparameter search space is described in Table 1 and Table 2 for SpineDL-Structure and SpineDL-Neuron respectively.

† These authors contributed equally to this work
* Corresponding authors

| **Hyperparameter** | **Search space** | **Best assignment** |
|---|---|---|
| Input image channels | *choice*[RGB (3), 2] | 2 |
| Patches | Extract patches from data with the minimum overlap | Extract patches from data with the minimum overlap |
| Patch size | *choice*[256 × 256, 512 × 512,<br>1024 × 1024] | 1024 × 1024 |
| Patch filtering | *choice*[none, by mean intensity] | none |
| Normalization | *choice*[uint8 division, scale_range,<br>zero_mean_unit_variance] | uint8 division |
| Percentile clipping | *choice*[none, (0.1,99.8)] | (0.1,99.8) |
| Data augmentation | *choice*[ Flips (horizontal + vertical),<br>Random rotations(-180,180),<br>Brightness(*choice*[(-0.2,0.2), (-0.1,0.1)<br>Contrast(*choice*[(-0.2,0.2), (-0.1,0.1)]),<br>Elastic transformations<br>Zoom(*choice*[(0.8,1.2), (0.9,1.1)]) ] | Flips (horizontal + vertical),<br>Random rotations(-180,180),<br>Brightness(-0.2,0.2),<br>Contrast(-0.2,0.2),<br>Elastic transformations |
| Epochs | *choice*[1500,1300,1000,800, 500] | 800 |
| Patience | *choice*[250,200,150,100,50] | 100 |
| Batch size | *choice*[2,3,4,6] | 2 |
| Loss type | BCE | BCE |
| Weights per class | *choice*[none, [0.74, 0.86, 1.71, 0.79,<br>1.89] ] | [0.74, 0.86, 1.71, 0.79, 1.89] |
| Optimizer | AdamW | AdamW |
| Learning rate | 1e-4 | 1e-4 |
| Warmup + cosine decay | Warmup epochs (*choice*[5,10]) | Warmup epochs (5) |
| Architecture | *choice*[U-Net, Residual U-Net,<br>Attention U-Net, HRNetV2-W48,<br>HRNetV2-W64] | HRNetV2-W48 |
| U-Net-like model feature maps and levels | *choice*[ $[16, 32, 64, 128, 256]$,<br>$[32, 64, 128, 256, 512]$ ] | - |
| U-Net-like model kernel size | *choice*[3,7] | - |
| Output prediction padding | *choice*[(64,64), (100,100), (200,200),<br>(400,400)] | (400,400) |
| Test-time augmentation | *choice*[yes, no] | Yes |

Table 1: **SpineDL-Structure hyperparameter search space.** More information about these keys can be found in BiaPy documentation.

| **Hyperparameter** | **Search space** | **Best assignment** |
|---|---|---|
| Input image channels | *choice*[RGB (3), 2] | 2 |
| Patches | Extract patches from data with the minimum overlap | Extract patches from data with the minimum overlap |
| Patch size | *choice*[256 × 256, 512 × 512] | 512 × 512 |
| Patch filtering | *choice*[none, by mean intensity] | By mean intensity |
| Normalization | *choice*[div, scale_range, zero_mean_unit_variance] | div |
| Percentile clipping | *choice*[none, (0.1,99.8)] | (0.1,99.8) |
| Data augmentation | *choice*[ Flips (horizontal + vertical), Random rotations(-180,180), Brightness(*choice*[(-0.2,0.2), (-0.1,0.1) Contrast(*choice*[(-0.2,0.2), (-0.1,0.1)]), Elastic transformations Zoom(*choice*[(0.8,1.2), (0.9,1.1)]) ] | Flips (horizontal + vertical), Random rotations(-180,180), Brightness(-0.2,0.2), Contrast(-0.2,0.2), Elastic transformations Zoom(0.8,1.2) |
| Epochs | *choice*[1500,1300,1000,800, 500] | 800 |
| Patience | *choice*[250,200,150,100,50] | 100 |
| Batch size | 6 | 6 |
| Loss type | BCE | BCE |
| Weights per channel | *choice*[none,[0.2,0.8],[1,2]] | [1,2] |
| Class rebalance within channel | *choice*[False, True] | True |
| Optimizer | AdamW | AdamW |
| Learning rate | 1e-4 | 1e-4 |
| Warmup + cosine decay | Warmup epochs (*choice*[5,10]) | Warmup epochs (5) |
| Architecture | *choice*[U-Net, Residual U-Net, Attent HRNetV2-W64 ] | HRNetV2-W48 |
| U-Net-like model feature maps and levels | *choice*[ [16, 32, 64, 128, 256], [32, 64, 128, 256, 512] ] | - |
| Output prediction padding | *choice*[(100,100), (50,50)] | (100,100) |
| Instance representation | *choice*[FC, FP] | FP |
| Seed creation channel | *choice*[FC, P] | P |
| Seed channel threshold | *choice*[auto, 0.5] | 0.5 |
| Topographic surface channel | F | F |
| Instance growth channel | F | F |
| Instance growth channel threshold | *choice*[auto, 0.9] | 0.9 |
| Filter instance seeds before watershed | *choice*[none, ¡10, ¡20, ¡65] | 10 |
| Filter instances (post-proc) | *choice*[none, ¡65, ¡100] | 100 |

Table 2: **SpineDL-Neuron hyperparameter search space.** More information about these keys can be found in BiaPy documentation.

### 2 Detailed Agreement Analysis

#### 2.1 Instance level metric for SpineDL-Neuron

We used common metrics to measure instance segmentation performance [1–4], which calculates a matching score between predicted and ground-truth (GT) instances based on a predefined IoU overlap. This instance-level approach simplifies performance assessment by classifying predictions as correct when the overlap exceeds a specified threshold. This evaluation strategy is consistent with those commonly employed in object detection tasks, where discrete instance identification is prioritized.

A predicted instance is considered a true positive (TP) if its IoU with a GT instance exceeds a predefined threshold. If this threshold is not met, the predicted instance is classified as a false positive (FP). Conversely, GT instances that do not match any predicted instance, or for which no predicted instance exceeds the IoU threshold, are considered false negatives (FN).

Using these statistics, the accuracy is computed as follows:

$$Accuracy = TP/(TP + FP + FN) \tag{1}$$

Following other state-of-the-art evaluations, and since neither the exact boundaries of spinal cord structures nor the full set of neurons present in each image is definitively known, we show metrics that require at least 50% IoU with the GT for a detection to be a TP.

#### 2.2 Neuron segmentation results

The precision-recall (PR) curve summarizes the balance between correctly detected neurons (recall) and the accuracy of those detections (precision) across different values of the seed-selection threshold applied to the center-of-mass (P) channel in the bottom-up instance segmentation process. Each point in the curve corresponds to a different threshold value, where lower thresholds allow more candidate neuron seeds to be generated, resulting in higher recall but lower precision, while higher thresholds restrict seed generation, producing fewer but more confident detections. The curve was computed on the validation dataset using the instance-level evaluation described above, in which predicted and ground-truth neurons were matched based on an IoU overlap of at least 50%, and TP, FP, and FN were derived accordingly. As is depicted in Figure 1, the analysis revealed that thresholds in the range of $0.2-0.4$ provided the most balanced trade-off between precision and recall, indicating optimal performance of the neuron instance segmentation pipeline within this range.

In addition to this parameter, the segmentation pipeline also includes a foreground constraint threshold used to limit the growth of regions during the marker-controlled watershed step. After qualitative inspection of the predictions, this constraint threshold was fixed at 0.9 to ensure well-defined neuron boundaries while avoiding over-segmentation.

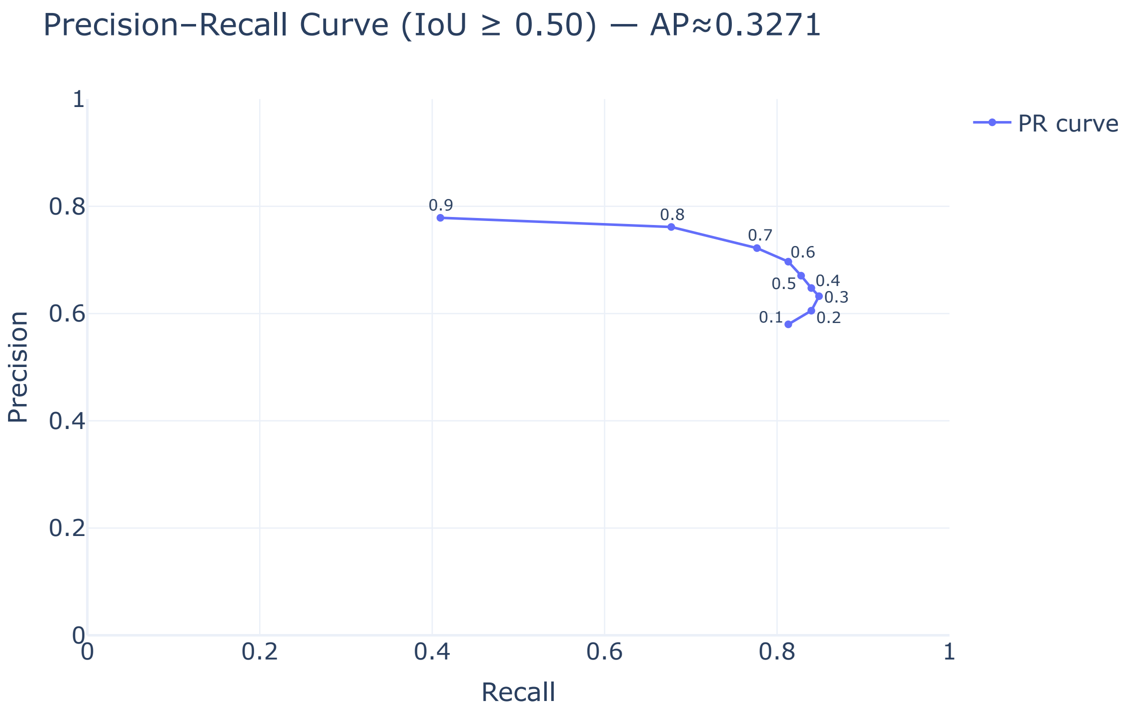


Fig. 1: **Precision-Recall (PR) curve for neuron instance segmentation using SpineDL-Neuron on the validation dataset.** Each point represents a different seed-selection threshold applied to the center-of-mass (P) channel, which defines candidate neuron seeds in the bottom-up segmentation process. A predicted neuron is considered a true positive when its IoU with a ground-truth instance exceeds 50%. The best balance between precision and recall is achieved for thresholds around $0.2 - 0.4$, reflecting the most reliable neuron identification performance.

### 3 Extra code availability

1. For SpineDL-Neuron, the evaluation between the predictions and the point annotations made by experts was performed using this script.
2. Foreground probabilities of CellSense were converted to instances using this script.
3. Script used for the figures:
   - Figure 4 and 5: script.
   - Figure 6: script.
   - Figure 7: script.
   - Figure S1: script.
